# Epithelial-Mesenchymal Transition is Associated with Altered Immune Composition and Cytotoxic Function in Triple-Negative Breast Cancer

**DOI:** 10.1101/2025.10.22.683714

**Authors:** Hanxu Lu, Meisam Bagheri, Fred W. Kolling, Lucas A. Salas, Xiaofeng Wang, Todd W. Miller, Diwakar R. Pattabiraman, Brock Christensen

**Affiliations:** Department of Molecular and Systems Biology, Geisel School of Medicine at Dartmouth, Lebanon, NH 03756, USA; Department of Epidemiology, Geisel School of Medicine at Dartmouth, Lebanon, NH 03756, USA; Dartmouth Cancer Center, Lebanon, NH 03756, USA; Department of Pharmacology & Toxicology, Medical College of Wisconsin, Milwaukee, WI, 53226, USA; Department of Pathology, Medical College of Wisconsin, Milwaukee, WI, 53226, USA

## Abstract

Epithelial-mesenchymal transition (EMT) is a dynamic process that contributes to breast cancer progression, metastasis, and therapy resistance. EMT also influences the tumor immune microenvironment, shaping immune cell infiltration and function. Although the immunological features of fully epithelial or fully mesenchymal tumor states have been characterized, the immune landscape associated with intermediate or partial EMT states remains poorly understood. To address this gap, we established five single-cell–derived clonal populations from the triple-negative 4T1 mouse mammary tumor cell line, representing a spectrum of EMT phenotypes, and analyzed tumors derived from these clones using single-cell RNA sequencing. We show that tumors derived from these clones retained their relative EMT states *in vivo*, and that EMT progression is associated with a graded reduction in tumor immunogenicity, accompanied by alterations in immune cell composition and function. Along the EMT spectrum, tumor cells exhibited progressive downregulation of major histocompatibility complex (MHC) class I and II gene expression, along with decreased infiltration of cytotoxic CD8+ T cells, and reduced expression of key effector and trafficking genes (*Gzmb*, *Ccr5*, *Cxcr6*). B cell composition also shifted, with decreased frequencies of IgG1-producing plasma cells and an enrichment of regulatory-like B cells. Natural killer (NK) cells similarly demonstrated progressive functional suppression, marked by reduced expression of cytotoxic molecules and the downregulation of key effector pathways. These findings reveal that EMT is associated with immune cell recruitment and function in a graded manner, providing a rationale for integrating EMT phenotyping into therapeutic strategies to overcome immune resistance in breast cancer.

## Introduction

Epithelial-mesenchymal transition (EMT) is a highly conserved biological process involved in embryonic development, wound healing, and cancer progression^1^. In the context of cancer, EMT facilitates the dissemination of tumor cells from the primary site by promoting the loss of epithelial features such as apical-basal polarity and cell-cell adhesion, and the acquisition of mesenchymal traits, including enhanced motility, invasiveness, and resistance to therapy^2,3^. Rather than occurring as a binary switch, EMT is increasingly recognized as a dynamic and plastic process, where cells transition through a spectrum of intermediate states that exhibit varying degrees of epithelial and mesenchymal characteristics^3–5^. These intermediate states, termed partial or hybrid EMT, exhibit enhanced tumor initiation capacity^6,7^, enabling collective migration and increasing metastatic risk^8^.

While extensive research has characterized the intrinsic properties of cancer cells undergoing EMT and their interactions with the tumor microenvironment (TME), much of this work has focused on comparisons between fully epithelial and fully mesenchymal states^9,10^. However, how immune cell infiltration and function are altered across the full EMT spectrum, particularly during partial EMT states, remains poorly understood.

To address this gap, we utilized five single-cell–derived clonal populations from the murine triple-negative 4T1 breast cancer cell line that are distributed along the EMT spectrum and retain these EMT states *in vivo*. Using single-cell RNA sequencing (scRNA-seq), we characterized the tumor microenvironment associated with each EMT state. Our data reveal progressive alterations in tumor cell antigen presentation and cytotoxic immune cell infiltration within the tumor microenvironment across the EMT spectrum, offering insights into how dynamic transitions, particularly partial EMT states, may contribute to immune evasion during tumor progression.

## Results

### Tumors derived from different EMT clones retained their original EMT states *in vivo*

To model the EMT spectrum in breast cancer, we generated five single-cell–derived clonal populations from the triple-negative (ER-/PR-/HER2-) 4T1 mouse mammary tumor cell line (Figure 1A) to represent the EMT spectrum. These included one epithelial-like (E) clone, three clones with intermediate states (EM1, EM2, EM3) representing partial EMT phenotypes, and one mesenchymal (M) clone. To rank clones along the EMT spectrum, they were assessed using expression of canonical EMT markers, *Cdh1* (E-cadherin), *Vim* (vimentin), *Snai1* (Snail), and *Twist1* (Twist1) (Figure 1B). To preserve the EMT characteristics of each clonal population, cells were maintained at low passage *in vitro* and discarded after a maximum of 10 passages. To assess *in vivo* tumorigenicity of each clonal population and the parental 4T1 line, they were orthotopically implanted into the fourth mammary fat pad of 9-week-old female BALB/c mice (n = 5 per group) (Figure 1C). Tumor growth kinetics revealed that the parental line and the intermediate clone EM2 exhibited the fastest growth, while the more epithelial-like clones (E and EM1) displayed moderate growth, and the mesenchymal-like clones (EM3 and M) showed the slowest growth (Figure 1D). These observations suggest a potential association between partial EMT states and enhanced tumorigenic capacity, as previously described by others^4^. To evaluate whether the EMT states were maintained *in vivo*, tumors were harvested upon reaching 1000 mm³ and processed into formalin-fixed paraffin-embedded (FFPE) blocks. Immunofluorescence staining for E-cadherin and vimentin was performed to visualize and quantify EMT marker expression (Figure 1E). The epithelial-like clone (E) derived tumors displayed the presence of cell-cell junctions, high E-cadherin expression, and low vimentin levels, consistent with an epithelial phenotype. In contrast, tumors from the more mesenchymal clones (EM2, EM3, and M) exhibited spindle-shaped morphology, reduced E-cadherin expression, and increased vimentin levels, characteristic of a mesenchymal state. Quantitative analysis of marker intensity revealed a progressive increase in the vimentin-to-E-cadherin ratio across the EMT spectrum (Figure 1F), with the epithelial clone (E) exhibiting the lowest ratio and the mesenchymal clone (M) the highest. This increasing trend was statistically significant, as determined by linear regression (p = 8.2×10⁻⁷). Notably, among the intermediate clones, EM2 exhibited the greatest heterogeneity in the vimentin/E-cadherin ratio, suggesting more phenotypic plasticity within this population. Together, these findings indicate that tumors derived from different EMT clones largely retained their original EMT states *in vivo*.

**Figure 1:**
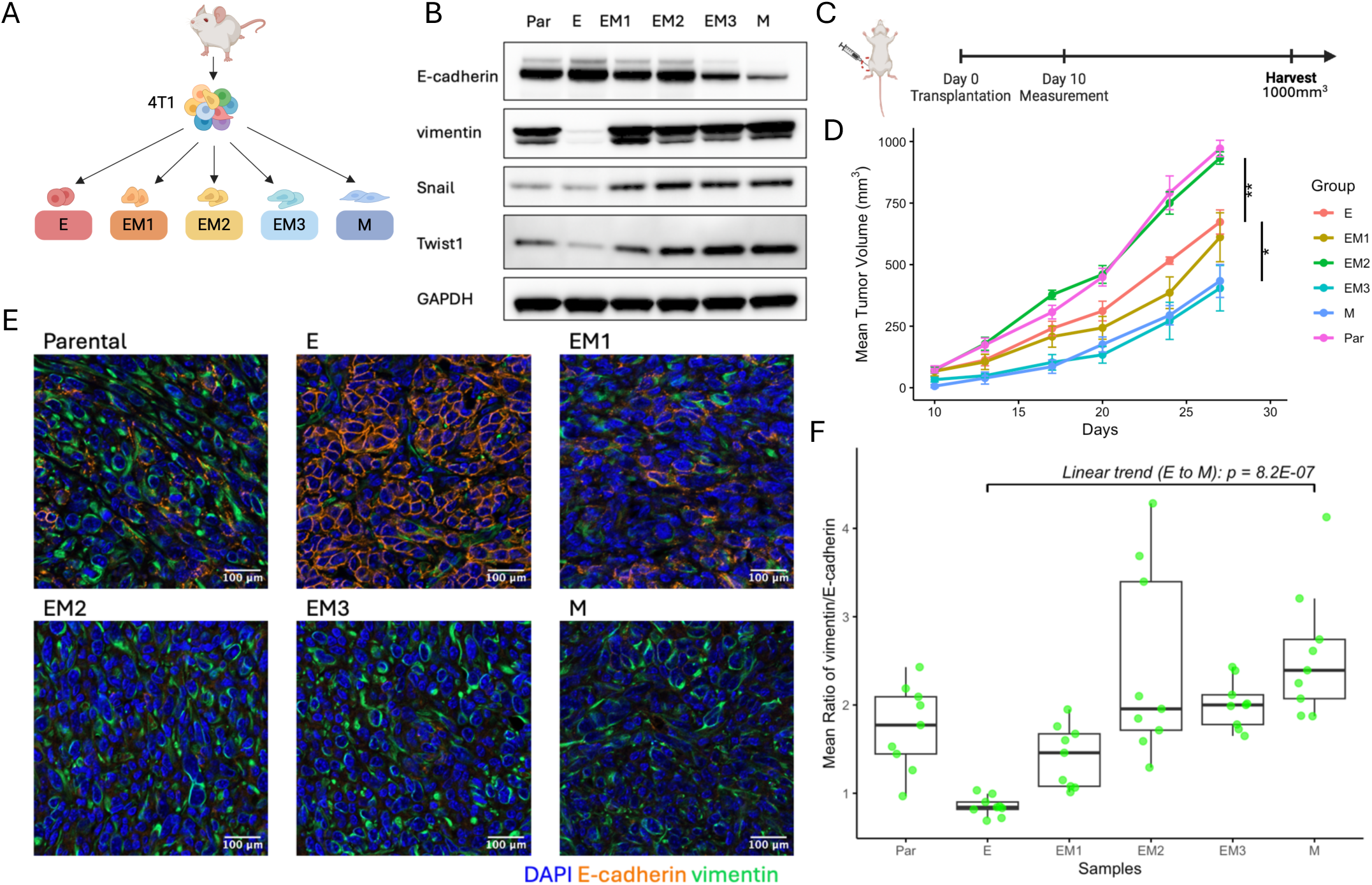
Establishment and characterization of a murine model to investigate the tumor EMT. **A** Schematic representation of the isolation process for single-cell–derived clonal populations at distinct EMT states from the 4T1 mouse breast cancer cell line. Clonal populations include an epithelial clone (E), three intermediate clones (EM1, EM2, EM3), and a mesenchymal (M) clone. **B** Immunoblot of canonical EMT markers, including E-cadherin, vimentin, Snail, and Twist1, to characterize the EMT states of isolated clones. GAPDH served as a loading control. **C** Experimental workflow: Six groups of mice (n = 5 per group) were orthotopically implanted with either EMT clones or parental 4T1 cells. **D** Tumor growth kinetics were measured twice per week, demonstrating distinct growth profiles associated with the EMT states of the injected clones. Data are shown up to the time point when the first tumor reached the endpoint volume. Mean tumor volumes (± standard error of the mean, SEM) are plotted over time. Statistical comparisons were performed using two-sided t-tests. Vertical lines and asterisks denote significant differences: p = 0.003 between E and EM2, and p = 0.022 between E and M. **E** Representative confocal immunofluorescence images of tumor sections derived from each EMT clone. Sections were co-stained for E-cadherin (orange), vimentin (green), and DAPI (blue) to visualize EMT marker expression *in vivo*. **F** Quantification of EMT marker expression across tumors. Each group includes three tumors, with three imaging fields analyzed per tumor. Data are shown as box plots of the normalized mean fluorescence intensity ratios of vimentin to E-cadherin. Fluorescence intensities were normalized to the DAPI signal. Linear regression analysis was performed to assess the trend in the vimentin-to-E-cadherin ratio along the EMT spectrum.

### Single-cell RNA sequencing identified distinct tumor and immune cell subpopulations in the tumor microenvironment

To investigate the relationship between EMT states and immune cell infiltration in the tumor microenvironment, we performed scRNA-seq on tumors derived from each clonally defined group. For each group, a representative tumor was selected based on sample quality and cell viability and processed to isolate both cells from total tumor tissue and matched CD45+ immune cells using magnetic-activated cell sorting (MACS) (Figure S1A). The scRNA-seq data from all individual cells across total tumor specimens were merged and integrated for Uniform Manifold Approximation and Projection (UMAP) analysis to visualize the cellular landscape within the tumor microenvironment (Figure 2A).

**Figure 2:**
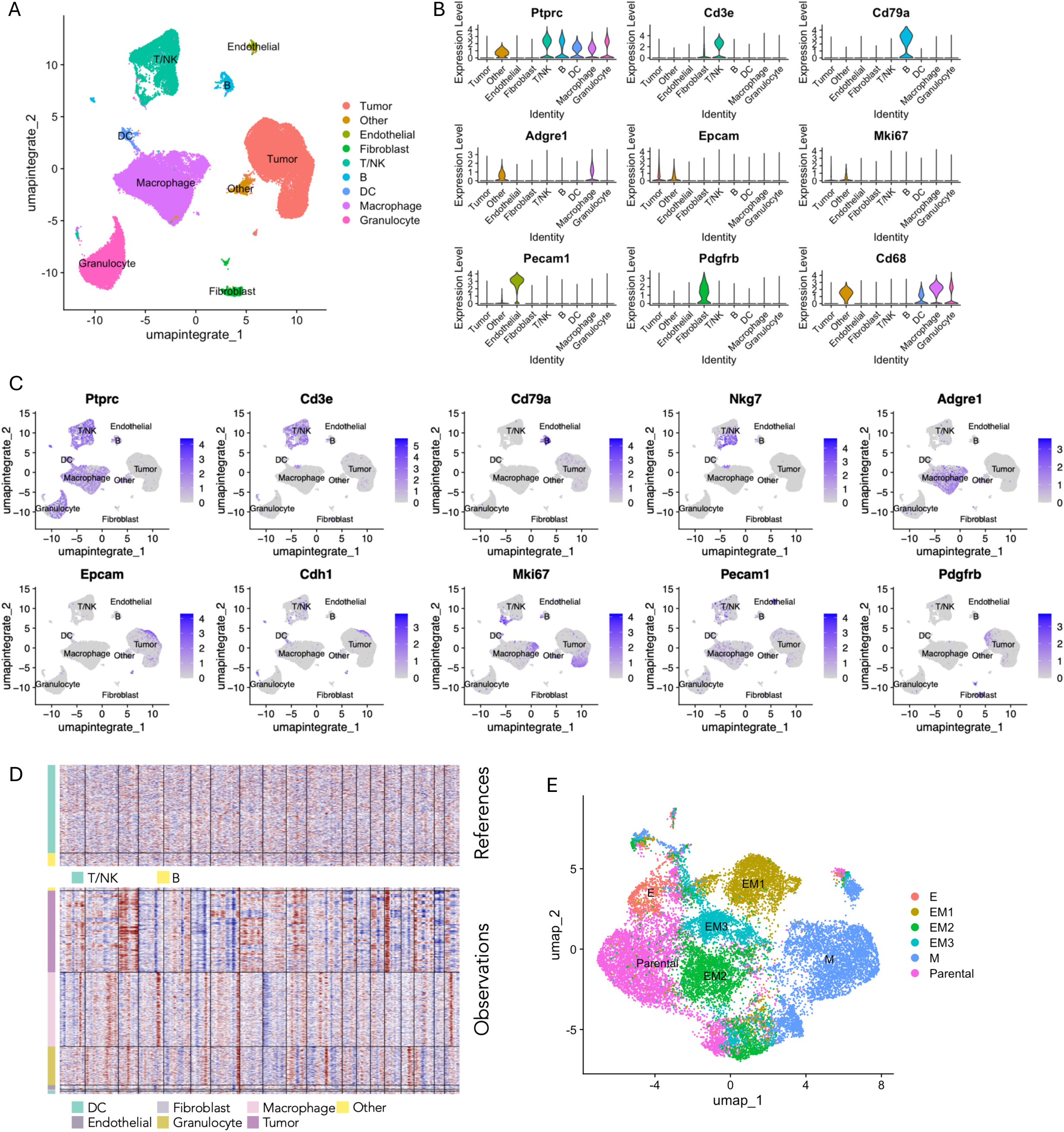
Single-cell transcriptomic profiling of the tumor microenvironment reveals cellular heterogeneity and EMT state distribution. **A** UMAP visualization of integrated scRNA-seq data from one representative tumor sample per group, illustrating the major cell populations within the tumor microenvironment. **B** Violin plots displaying the expression levels of key marker genes across each identified cell cluster. **C** Feature plots demonstrating the expression of key marker genes specific to the major cell types within the tumor microenvironment **D** Copy number variation (CNV) profile generated to confirm the identification of tumor cells. The x-axis represents genomic regions ordered by chromosome 1 to 19. The y-axis shows individual cells, with reference cells (T/NK and B cell clusters) displayed in the top panel and observed cells in the bottom panel. Red indicates relative copy number gain, and blue indicates relative loss. **E** UMAP visualization of re-clustered tumor cells, colored by sample of origin.

Based on transcriptional similarity observed in UMAP space, distinct cell populations were manually identified and annotated using canonical lineage markers: tumor cells (*Epcam+ Cdh1+*), T/NK cells (*Cd3d+*, *Nkg7+*), B cells (*Cd79a+*), macrophages (*Adgre1+*, *Cd68+*), endothelial cells (*Pecam1+*), and fibroblasts (*Pdgfrb+*) (Figure 2B-C, Figure S1C). Cluster identities were confirmed using SingleR, an unbiased automated clustering approach^11^ using the Immunological Genome Project^12^ (ImmGen) reference database (Figure S1B). To validate the identification of tumor cells, we conducted copy number variation (CNV) analysis using a computational approach that infers large-scale chromosomal alterations from scRNA-seq data^13,14^. Using T/NK and B cell clusters as reference populations, the inferred CNV profiles revealed chromosomal instability in tumor cells, characterized by widespread and irregular amplifications and deletions (Figure 2D), distinguishing malignant cells from stromal and immune components.

Tumor cells were stratified and visualized using UMAP, revealing distinct clusters corresponding to cells derived from different EMT clones or the parental population (Figure 2E, Figure S1E). Each EMT state formed a relatively discrete cluster in the UMAP space, suggesting that the core transcriptional features defining each EMT state are robust enough to drive the segregation of tumor cells by EMT states. Parental cells also formed a distinct cluster, largely separate from EMT-derived populations, reflecting inter-tumor heterogeneity. Notably, while most clusters were enriched for cells from a single clonal origin, some clusters contained cells from multiple clones, indicating that despite the transcriptional stability of EMT states, there is a degree of phenotypic plasticity and intra-tumor heterogeneity within individual tumors.

### Gene expression profiling of tumors from different clonal EMT states reveals epithelial and mesenchymal markers and immune evasion mechanisms

Following tumor cell identification, we isolated the tumor cluster from the scRNA-seq dataset and assessed EMT states by calculating EMT scores at the single-cell level using the 76-gene signature (76GS) method^15,16^. This approach calculates EMT scores based on the correlation between the expression levels of pre-determined EMT-related genes (Table S1) and *Cdh1* (E-cadherin), a canonical epithelial marker^15^. Higher EMT scores reflect an epithelial-like phenotype, whereas lower scores correspond to a mesenchymal state. Analysis of our scRNA-seq data revealed that tumor cells derived from the most epithelial-like clone (E) exhibited the highest EMT scores, consistent with the retention of epithelial characteristics as observed by immunofluorescence staining (Figure 3A, Figure 1E). In contrast, tumors originating from more mesenchymal-like clones displayed progressively reduced EMT scores, reflecting their mesenchymal nature (Figure 3A). These transcriptome-based EMT scores are consistent with immunofluorescence-based assessments, supporting the maintenance of clone-specific EMT states *in vivo* and providing a complementary, high-resolution view of EMT heterogeneity within tumors.

**Figure 3:**
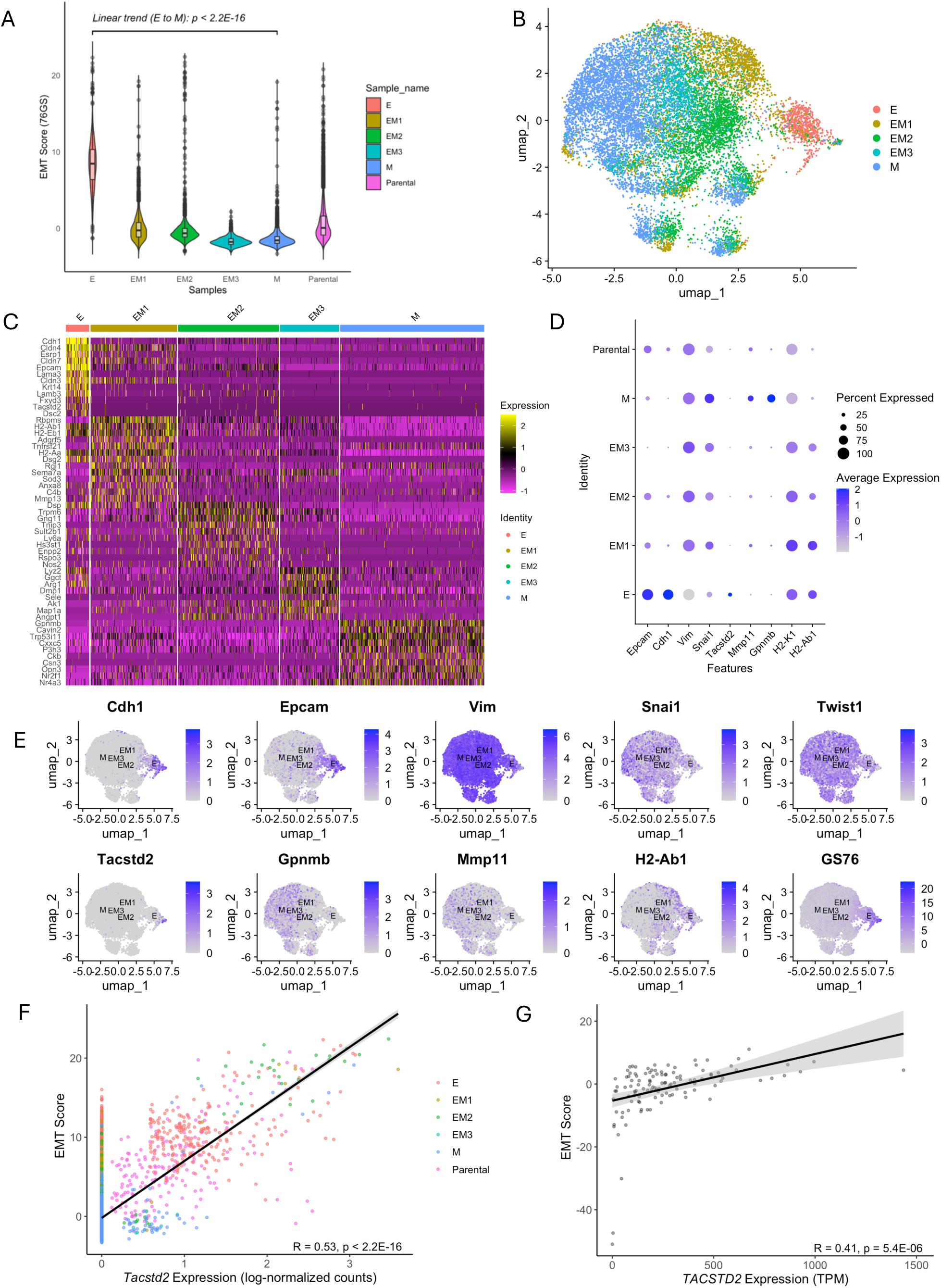
Characterization of EMT score profiling and differential gene expression across tumor cell clusters from distinct EMT clones. **A** EMT scores are calculated at the single-cell level using the 76-gene signature (76GS) method after subsetting tumor cells from the five EMT clones and the parental 4T1 line. Linear regression analysis was performed to assess the trend in EMT scores from E to M. **B** UMAP visualization of scRNA-seq data showing the isolated tumor cell cluster from the five EMT clones using EMT-related genes, excluding cells from the parental line. **C** Heatmap displaying the most variably expressed genes (log₂ fold change > 1.5) across different tumor cell populations. **D** Dot plot showing the average expression levels of selected epithelial and mesenchymal markers across tumor samples. Dot size represents the percentage of cells expressing each gene, color intensity indicates the average expression level. **E** UMAP feature plots showing 76GS EMT scores and expression of EMT-associated genes. **F** Scatter plot showing the correlation between *Tacstd2* expression and EMT scores in tumor cells derived from EMT clones (R = 0.53, p < 2.2 × 10^-16^), based on scRNA-seq data. Each point represents an individual cell, colored by tumor clone identity. **G** Correlation between *TACSTD2* expression and EMT scores in ER-/PR-/HER2- breast cancer patients from the TCGA-BRCA cohort using bulk RNA-seq data (R = 0.41, p = 5.4 × 10^−6^). Each point represents a patient primary tumor sample.

To validate the EMT states observed with the 76GS method, we applied two additional EMT scoring approaches reported in previous studies: a Kolmogorov–Smirnov (KS) statistic-based approach^17^ and a multiple linear regression (MLR)-based method^18^. Both scoring metrics were adjusted to align with the directional interpretation of the 76GS score, wherein higher scores indicate a more epithelial phenotype (Figure S2C). High correlation coefficients between EMT scores derived from the KS, MLR, and 76GS methods demonstrated strong concordance among these approaches (Figure S2D), reinforcing the robustness and reliability of EMT state classification across methodologies. Furthermore, when compared to EMT scores from 81 murine mammary cancer cell lines from the TISMO database^19^, tumor cells derived from the intermediate EMT clones and the mesenchymal EMT clone occupied an intermediate position along the broader EMT spectrum (Figure S2E). This trend was consistent across all three EMT scoring methods, suggesting that although tumors derived from distinct EMT clones retained their relative EMT hierarchy upon tumor formation *in vivo*, they may adopt a hybrid or partial EMT phenotype over time. The observed phenotypic plasticity may facilitate dynamic transitions along the EMT spectrum, contributing to intra-tumor heterogeneity.

To further investigate transcriptomic variation across EMT clone–derived tumors, we isolated the tumor cell cluster, excluded parental 4T1 cells, and analyzed the most variably expressed genes across samples and performed dimensionality reduction again using a set of EMT-related genes (Table S1). UMAP analysis revealed a spectrum of five distinct transcriptional states, representing a gradual transition from epithelial to mesenchymal phenotypes (Figure 3B). The intermediate states formed a gradient between the two extremes, supporting the existence of partial EMT states rather than a binary switch. Tumors derived from the epithelial-like (E) clone exhibited high expression of key epithelial markers, including *Cdh1*, *Epcam*, *Cldn4, and Cldn7*, along with low expression of mesenchymal markers such as *Vim* and *Snai1* (Figure 3C-E).

Notably, tumors from the most mesenchymal-like (M) clone exhibited elevated expression of *Gpnmb* (Figure 3C), a gene frequently co-upregulated with matrix metalloproteinase (MMP) genes and associated with increased cancer metastasis^20,21^. *Gpnmb* has also been implicated in promoting immune evasion, as it can attenuate T cell activation^21,22^. Comparison of the average expression of a selected panel of genes across samples revealed that *Gpnmb* and *Mmp11* were co-upregulated in tumor cells from the most mesenchymal-like (M) clone, further supporting the association of these genes with a mesenchymal-like phenotype (Figure 3D). Additionally, a progressive downregulation of major histocompatibility complex (MHC) class I and II molecules was observed in tumors with increasing mesenchymal features (Figure 3D and Figure S1D). This downregulation of antigen presentation machinery may impair immune recognition, thereby limiting T cell recruitment and reducing immune surveillance. Collectively, these observations suggest that the EMT process may facilitate immune evasion by altering antigen presentation pathways, impairing T cell recruitment, and promoting an immunosuppressive tumor microenvironment.

Among the top differentially expressed genes in the epithelial-like (E) clone, *Tacstd2*, which encodes TROP2, was also one of the most highly upregulated genes (Figure 3C-E). *Tacstd2* expression was positively correlated with EMT scores in both our mouse tumor scRNA-seq dataset (Figure 3F) and human triple-negative breast cancer (TNBC) samples from the TCGA-BRCA cohort, based on bulk RNA-seq data (Figure 3G). This suggests that TROP2 expression is associated with EMT states in both murine and human TNBC. TROP2 has been widely recognized as a therapeutic target and is commonly linked to E-cadherin expression and a stable epithelial phenotype^23,24^. Our results indicate that TROP2 expression may vary along the EMT spectrum, raising the possibility that EMT states could affect sensitivity to TROP2-targeted antibody–drug conjugates (ADCs) such as sacituzumab govitecan^25^. Integrating EMT states with TROP2 expression may therefore improve therapeutic stratification and enhance prediction of ADC response in TNBC.

### Distinct EMT states are associated with changes in T cell composition and cytotoxic gene expression

To examine the relationship between tumor EMT states and immune cell infiltration, we analyzed CD45+ immune cells isolated by MACS from the same six tumors (five EMT clone– derived and one parental) using scRNA-seq (Figure S1A). Samples were merged and integrated, and UMAP visualization revealed distinct immune lineages (Figure 4A, Figure S3B). Cluster identities were validated using the ImmGen reference database (Figure S3A). The T cell population was identified by *Cd3e* expression and divided into seven transcriptionally distinct subclusters (Figure 4B-C). These included a cytotoxic CD8+ T cell population characterized by high expression of *Gzmb* and *Ccr7* (Gzmb+ CD8+ T), a precursor-exhausted CD8+ T cell cluster co-expressing *Tcf7, Lef1* and *Pdcd1^26–28^*(Lef1+ CD8+ T), and a regulatory T cell (Treg) population expressing *Cd4* and *Foxp3* (Foxp3+ CD4+ T) (Figure 4D). Additional subpopulations included Cd44-Cd4+ T cells, Cd44+ Cd4+ T cells, proliferating Mki67+ T cells, and Trdc+ double-negative (DN) T cells, which likely correspond to gamma-delta (γδ) T cells^29^.

**Figure 4:**
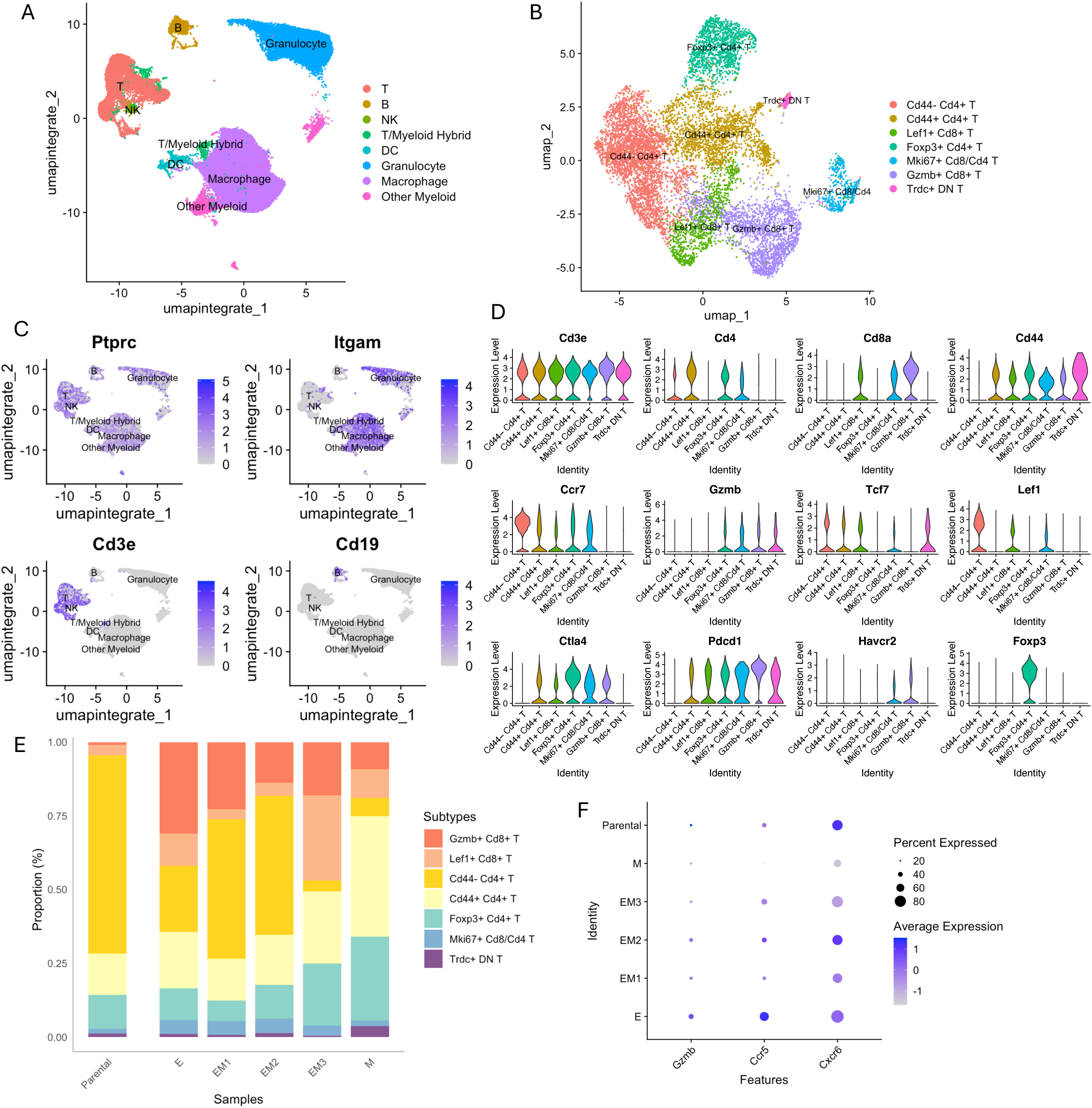
Impact of EMT states on T cell subtype composition and gene expression in the tumor microenvironment. **A** UMAP plot merged CD45+ immune cell–enriched samples from six tumor samples to identify major immune cell lineages. **B** UMAP visualization of T cell subclusters. **C** Feature plots showing expression of representative markers (*Ptprc*, *Itgam*, *Cd3e*, *Cd19*) to distinguish T cells, macrophages, and B cell lineages in CD45+ cells. **D** Violin plots showing expression of canonical genes to identify T cell subtypes. **E** Stacked bar plot showing the proportions of T cell subtypes across different samples. **F** Dot plot showing average expression of *Gzmb*, *Ccr5*, and *Cxcr6* in cytotoxic CD8+ T cells across different samples.

When comparing the proportions of T cell subtypes across clonally derived tumors with varying EMT states, the abundance of T cells progressively declined from epithelial-like to mesenchymal-like tumors, with the most pronounced reduction in the proportion of cytotoxic CD8+ T cells (Gzmb+ CD8+ T) (Figure 4E, Figure S3D). In addition, the proportion of Tregs exhibited a progressive increase along the EMT spectrum. These observations indicate that mesenchymal-like tumors are associated with a less immunogenic microenvironment. To explore this further, we analyzed gene expression in cytotoxic CD8+ T cells and found that cells from more mesenchymal-like tumors exhibited lower expression of *Gzmb*, *Ccr5*, and *Cxcr6* compared to those from more epithelial-like tumors (Figure 4F). Given that CCR5 and CXCR6 are involved in T cell migration and retention within tumor tissues^30–33^, and that granzyme B (*Gzmb*) is a key effector molecule for cytotoxic function^34^, these findings suggest that as tumors progress toward a mesenchymal-like EMT state, they impair both the recruitment and functional activity of cytotoxic T cells, contributing to a less immunogenic tumor microenvironment.

### Tumor EMT associates with shifts in B cell subpopulations and progressive suppression of NK cell cytotoxic function in the tumor microenvironment

To assess how tumor EMT states relate to B cell dynamics in the tumor microenvironment, we identified B cells based on *Cd19* expression in immune-enriched scRNA-seq data (Figure 4C) and visualized the data using UMAP, which revealed four transcriptionally distinct B cell subpopulations: Cd44-MHCIIhi B cells, plasma cells, Cd25+ Cd69+ B cells, and Cd44+ Cd69+ B cells (Figure 5A-B). Similar to the T cell compartment, the relative proportions of these B cell subpopulations also varied across tumors with different EMT states, indicating a dynamic shift in the immune cell composition as tumors progress from epithelial-like to mesenchymal-like phenotypes.

**Figure 5:**
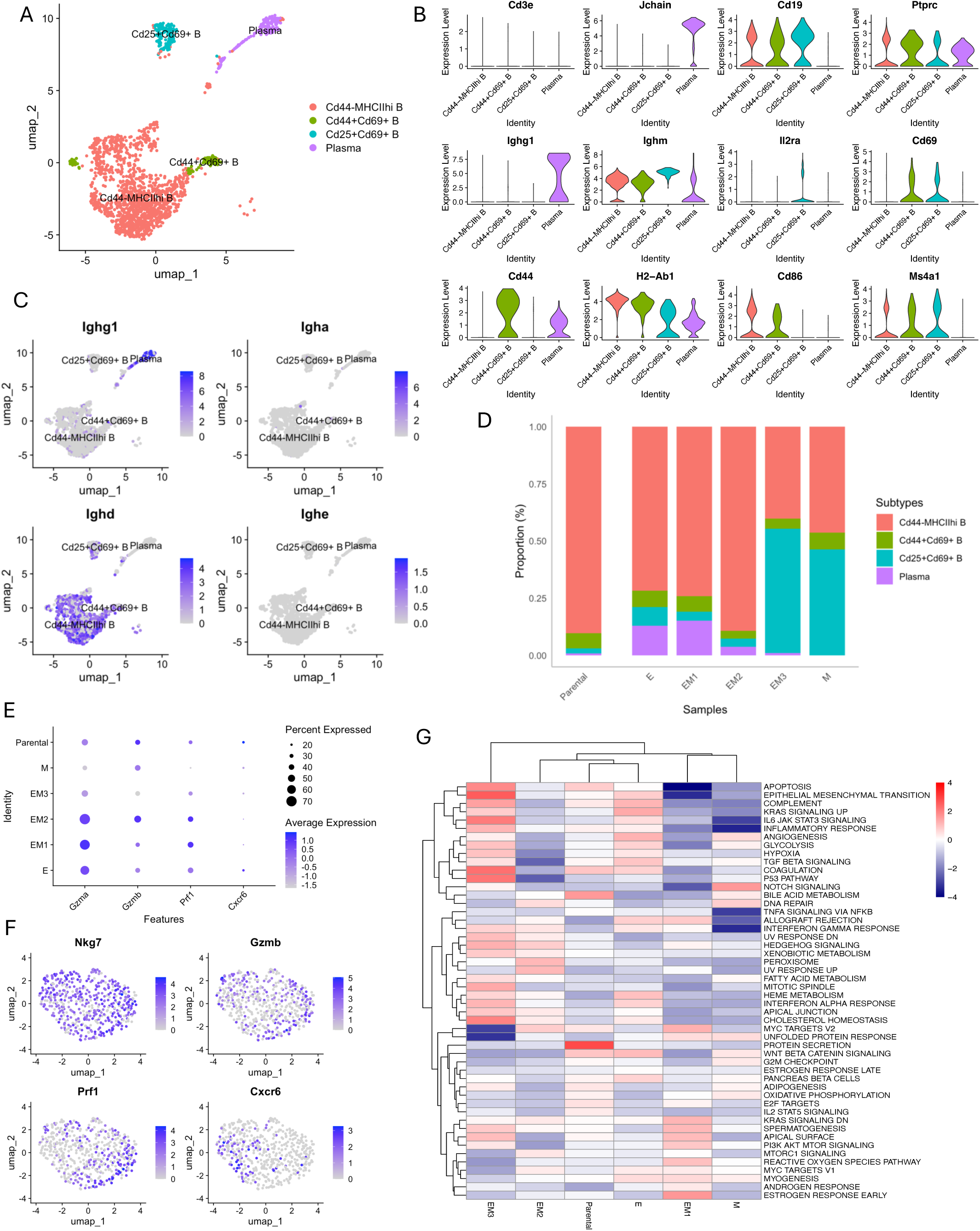
EMT-associated alterations in B cell and NK cell populations in the tumor microenvironment. **A** UMAP plot of B cells derived from CD45+ cell enriched samples across tumors with different EMT states, four distinct B cell subpopulations were identified: Cd44-MHCIIhi B cells, plasma cells, Cd25+Cd69+ B cells, and Cd44+Cd69+ B cells. **B** Violin plots showing expression of representative marker genes across B cell subtypes. **C** Feature plots displaying expression of immunoglobulin isotypes (*Ighg1, Igha, Ighd, Ighe*) in B cells. **D** Stacked bar plot illustrating the relative proportions of B cell subtypes in tumors with different EMT states. **E** Dot plot showing average expression of *Gzma, Gzmb, Prf1, and Cxcr6* in NK cells across different samples. **F** Feature plots displaying expression of *Nkg7, Gzmb, Pfr1*, and *Cxcr6* in NK cells. **G** Heatmap of Z-scores representing pathway enrichment in NK cells based on scRNA-seq data. Hallmark gene sets from the MSigDB database were used for gene set enrichment analysis (GSEA), with enrichment calculated using the Wilcoxon–Mann–Whitney. Z-scores reflect the relative enrichment of each pathway across samples.

In tumors that retained a more epithelial-like phenotype (E, EM1), plasma cells were enriched (Figure 5D) and predominantly expressed *Ighg1* (Figure 5C). As IgG antibodies, particularly IgG1, have been linked to anti-tumor B cell responses^35^, this enrichment suggests the presence of immune-reactive plasma cells in a more epithelial-like tumor microenvironment. In contrast, plasma cell proportions were markedly reduced or absent in more mesenchymal-like tumors (EM3, M), this decline may lead to a decrease in IgG1-mediated anti-tumor immunity, reflecting a more immunosuppressive tumor microenvironment.

Notably, Cd25+ Cd69+ B cells, which mark regulatory B cells (Bregs) in 4T1 tumors^36^, were enriched in more mesenchymal-like tumors (EM3 and M) (Figure 5D). Bregs are known to promote immune suppression through mechanisms such as T cell exhaustion and the inhibition of effective anti-tumor responses^37^. Their enrichment in more mesenchymal-like tumors suggests an immunosuppressive shift in the B cell compartment, potentially complementing the observed reduction in cytotoxic CD8+ T cell infiltration. These coordinated alterations, together with the downregulation of MHC molecules in tumor cells, highlight that EMT-associated immune evasion is mediated by a convergence of multiple pathways that collectively reshape the immune landscape.

To evaluate natural killer (NK) cell function across the EMT spectrum, we then isolated NK cells from CD45+ scRNA-seq samples (Figure 5F) and assessed both cytotoxic gene expression and pathway activity. A dot plot revealed a progressive decrease in both expression levels and the proportion of NK cells expressing canonical cytotoxic and trafficking-associated genes (*Gzma*, *Gzmb*, *Prf1*, *Cxcr6*) from epithelial to mesenchymal tumors (Figure 5E). This trend suggests a gradual impairment of NK cell cytolytic capacity as EMT progresses.

Pathway enrichment analysis further revealed that NK cells from mesenchymal-like tumors exhibited downregulation of key immune effector programs, including TNFα/NFκB signaling, IFN-γ response, allograft rejection, and IL6-JAK-STAT3 signaling, indicating attenuated activation and cytokine responsiveness (Figure 5G). These findings suggest that EMT contributes to the establishment of a tumor microenvironment that suppresses NK cell effector function and impairs anti-tumor immunity.

## Discussion

Epithelial-mesenchymal transition (EMT) is a well-established driver of cancer progression, metastatic dissemination, and therapy resistance^1,2,38^. However, its role in modulating tumor-immune interactions remains incompletely understood, particularly in the context of partial EMT states. In this study, we used single-cell–derived clonal populations representing distinct EMT states from the 4T1 murine triple-negative breast cancer model to investigate how tumor EMT progression influences the immune landscape of the tumor microenvironment. Our findings demonstrate that EMT state is associated with profound shifts in both tumor cell–intrinsic gene expression and the composition and function of infiltrating immune cells including T, B, and NK cell lineages.

The five EMT clones exhibited transcriptionally distinct profiles, as evidenced by their separation into discrete clusters in the scRNA-seq analysis. Although some EMT clone–derived tumors occupied a more intermediate position along the EMT spectrum compared to the broader panel of mammary tumor cell lines in the TISMO dataset, they retained their relative EMT hierarchy established *in vitro*. This was supported by consistent expression patterns of epithelial and mesenchymal markers observed through immunofluorescence and EMT scoring derived from scRNA-seq data. The preservation of these EMT states *in vivo* enabled the robust assessment of how defined EMT phenotypes influence immune cell infiltration within a controlled genetic background. Tumors with more mesenchymal-like features showed reduced expression of MHC class I and II genes, aligning with prior studies linking EMT to impaired antigen presentation and reduced immunogenicity^39^. This EMT-associated downregulation of antigen presentation machinery may contribute to immune evasion by limiting recognition and activation of cytotoxic immune cells within the tumor microenvironment.

In breast cancer, tumor cells at the fully mesenchymal end of the EMT spectrum have been associated with increased infiltration of immunosuppressive populations such as Tregs, M2 macrophages, and exhausted CD8+ T cells^39^, which collectively impair immune function and promote tumor progression ^40,41^. While these associations are well established at the extremes of EMT, our study provides new evidence that immune suppression develops progressively across the EMT spectrum. We observed a gradual reduction in cytotoxic CD8+ T cell infiltration and effector gene expression (*Gzmb, Ccr5, Cxcr6*), along with a corresponding increase in Tregs, from epithelial to mesenchymal states. These changes align with tumor-intrinsic alterations in MHC molecule expression and support a model in which EMT-associated immune infiltration is a continuous process rather than a discrete event.

Our study reveals previously unrecognized alterations in B cell and NK cell compartments across the EMT spectrum. While IgG-producing plasma cells have been associated with improved overall survival^42^, our findings demonstrate a progressive decline in their abundance from epithelial to mesenchymal states, accompanied by an increase in Cd25+ Cd69+ regulatory B cells. At the same time, NK cells exhibited a gradual loss of cytotoxic gene expression and downregulation of key effector pathways, including TNFα/NFκB, IFN-γ response, and IL6-JAK-STAT3 signaling, indicating a shift toward a less cytotoxic, more immunosuppressed state.

While NF-κB has been shown to regulate immune cell activation and *Ifng* transcription in NK cells^43^, our data indicate that EMT suppresses these signaling axes within NK cells, potentially impairing innate immune surveillance. Together, these findings suggest that as tumors progress toward a mesenchymal-like EMT state, they promote immune evasion by coordinately modulating both tumor-intrinsic features and the surrounding immune landscape.

Interestingly, we identified *Tacstd2* (TROP2), a clinically relevant transmembrane glycoprotein^44^, as one of the most highly expressed genes in tumor cells derived from the epithelial-like (E) clone. TROP2 expression was positively correlated with EMT scores in both our single-cell dataset and human TNBC samples from the TCGA-BRCA cohort. As the therapeutic target of Sacituzumab govitecan, an FDA-approved antibody–drug conjugate (ADC) for metastatic TNBC^45^, TROP2 represents a key biomarker for targeted therapy. However, our data indicate that TROP2 expression declines during EMT progression, including in intermediate or hybrid states, suggesting that EMT plasticity may reduce target availability and potentially limit the effectiveness of TROP2-directed therapies, in addition to rendering cells more resistant to conventional chemotherapy. Accordingly, assessing both EMT states and TROP2 expression levels may provide a more refined approach for identifying TNBC patients most likely to benefit from ADC therapy.

While our EMT clonal model provides a high level of internal consistency and enables the controlled study of defined EMT states within a uniform genetic background, it is derived from a single murine mammary carcinoma cell line. As such, the extent to which these findings can be generalized to other tumor types, species, or more genetically diverse models remains to be determined. In human tumors, both intra-tumor and inter-tumor heterogeneity, as well as the presence of mixed or partial EMT phenotypes, may limit the ability of EMT classification alone to accurately predict immune cell composition or functional immune responses within the tumor microenvironment.

Moreover, although our single-cell analysis reveals associations between EMT states and immune remodeling within the tumor microenvironment, causal relationships remain to be established and will require functional validation using inducible EMT systems or genetic manipulation of key EMT regulators. The bidirectional relationship, in which the tumor microenvironment can induce EMT and EMT in turn modulates immune composition, highlights the dynamic interplay between tumor plasticity and immune surveillance. These findings underscore the need for experimental models that more accurately reflect this complexity to uncover the mechanisms underlying EMT-driven immune modulation.

In conclusion, our study demonstrates that the EMT state is a critical determinant of immune cell composition and function within the triple negative breast tumor microenvironment. By shaping both the antigenic visibility of tumor cells and the recruitment and activation of immune effectors, EMT contributes to immune evasion in a state-dependent manner. These findings highlight the importance of accounting for EMT states in the design and implementation of immunotherapies, and they may provide a rationale for combination strategies that target both EMT and immune checkpoints in aggressive, mesenchymal-like tumors.

## Methods

### Cell Culture

The mouse 4T1 cell line (ATCC Cat# CRL-2539, RRID:CVCL_0125) was maintained in 1× RPMI 1460 (Corning, Cat# 10-040-CV) medium with 10% fetal bovine serum (FBS) (Gibco, Cat# 10438-026) and 1% penicillin-streptomycin (Corning Cat# 30-002-CI). The cells were incubated at 37°C in a 5% CO2-air atmosphere with constant humidity. Trypsin–EDTA (Corning Cat# 25-053-Cl) was used for passaging, serum-free cell freezing medium (Bambanker Cat# 302-14681) was used for freezing. Passage numbers were recorded for all cell lines, and cells were discarded after a maximum of 10 passages to preserve their EMT characters.

### EMT Clone Generation

Fluorescence-activated cell sorting (FACS) was performed on parental 4T1 (ATCC Cat# CRL-2539, RRID:CVCL_0125) cells using a FACSAria III Cell Sorter (BD Biosciences, RRID:SCR_016695). Cells were stained with APC anti-mouse EpCAM Antibody (BioLegend Cat# 118214, RRID:AB_1134102, 1:100) followed by 4′,6-Diamidine-2′-phenylindole dihydrochloride (DAPI, Sigma-Aldrich Cat# 10236276001). Cells were gated based on EpCAM expression levels into three groups, sorted into collection tubes, and immediately seeded into 6-well plates. Once 70% confluent, cells were trypsinized and plated at a density of one cell per well in 96-well plates. Single-cell–derived clones were expanded and screened for EMT characteristics.

### Immunoblotting

Cell lysates were harvested directly from culture plates using cell scrapers in 1× RIPA buffer (Sigma-Aldrich Cat# 20-188) with protease and phosphatase inhibitors (Thermo Fisher Scientific Cat# 78447). Protein concentrations were measured using the Bradford assay, and 40 μg of total protein per lane was loaded onto a NuPAGE Bis-Tris gel (Thermo Fisher Scientific). Proteins were transferred to nitrocellulose membranes using the iBlot semidry transfer system (Thermo Fisher Scientific), then blocked in blocking buffer with TBST containing 2.5% milk and 2.5% bovine serum albumin (Sigma-Aldrich Cat# A9647). Membranes were incubated overnight with the following primary antibodies: E-cadherin (BD Biosciences Cat# 610182, RRID:AB_397581, 1:1000), Vimentin (Cell Signaling Technology Cat# 5741, RRID:AB_10695459, 1:2000), Snail (Cell Signaling Technology Cat# 3879, RRID:AB_2255011, 1:1000), Twist1/2 (Abcam Cat# ab50887, RRID:AB_883294, 1:50), and GAPDH (Cell Signaling Technology Cat# 2118, RRID:AB_561053, 1:1000). HRP-conjugated goat anti-mouse IgG (H+L) (Thermo Fisher Scientific Cat# 31431, RRID:AB_10960845) and goat anti-rabbit IgG (H+L) (Thermo Fisher Scientific Cat# 31466, RRID:AB_10960844) secondary antibodies were applied at a 1:10000 dilution for 1 hour at room temperature in the blocking buffer. Blots were developed and imaged using the Bio-Rad ChemiDoc MP Imaging System (RRID:SCR_019037).

### *In Vivo* Tumor Implantation and Monitoring

Cells were digested and resuspended in Opti-MEM (Thermo Fisher Scientific Cat# 11058021) with 20% Matrigel (VWR Cat# 47743-706), and 100000 cells in 50 μL were implanted into the fourth mammary fat pad of female BALB/c mice (RRID:IMSR_JAX:000651) under isoflurane anesthesia. Tumor volume was measured twice weekly using calipers, and volume calculated with the formula ½ (length × width²). Tumors were removed by a live surgery at the time of tumor burden (total tumor volume of 1000 mm3). Mice were euthanized at endpoint, and tumors, lungs, and livers were harvested and fixed overnight in 10% neutral-buffered formalin for paraffin embedding.

### Tumor Dissociation

Freshly harvested tumors were digested in RPMI medium containing 2 mg/mL collagenase A (Sigma-Aldrich Cat# 10103586001) and 100 U/mL hyaluronidase (Fisher Scientific Cat# ICN10074091) at 37°C with rotation for 2 hours. The resulting suspension was sequentially filtered through 70 μm (Corning Cat# 431751) and 40 μm (Corning Cat# 431750) cell strainers. Red blood cells were lysed using 1× RBC lysis buffer (prepared from NH4Cl 8.02 g, NaHCO3 0.84 g, EDTA 0.37 g in 100 mL water for 10× stock). Tumor-infiltrating immune cells were enriched using mouse CD45 (TIL) MicroBeads (Miltenyi Biotec Cat# 130-110-618, RRID:AB_3224404) to separate CD45+ and CD45- fractions.

### Tissue Immunofluorescence Staining

Formalin-fixed, paraffin-embedded (FFPE) tissue blocks were sectioned at 4 μm thickness. Slides were dewaxed using xylene (VWR Cat# EMD-XX0055-5) and rehydrated through an ethanol gradient. Antigen retrieval was performed in a pressure cooker using Leica BOND Epitope Retrieval Solution 1 (Leica Cat# AR9961). Permeabilization was carried out using 1× TBS containing 0.5% Triton X-100 (Sigma-Aldrich Cat# 648466). Slides were blocked with 1% bovine serum albumin (BSA, Sigma-Aldrich Cat# A9647) and 3% normal goat serum (Thermo Fisher Scientific Cat# 50062Z) in 1× PBS (Thermo Fisher Scientific Cat# 70011069). Primary antibodies against E-cadherin (BD Biosciences Cat# 610182, RRID:AB_397581, 1:200) and Vimentin (Cell Signaling Technology Cat# 5741, RRID:AB_10695459, 1:200) were applied overnight at 4°C in blocking buffer. Secondary antibodies (anti-rabbit, Thermo Fisher Scientific Cat# A-11034, RRID:AB_2576217, 1:2000) (anti-mouse, Thermo Fisher Scientific Cat# A-21422, RRID:AB_2535844, 1:2000) and DAPI (DAPI, Sigma-Aldrich Cat# 10236276001) in blocking buffer were used for 1 hour incubation. Slides were mounted using ProLong Diamond Antifade Mountant (Invitrogen Cat# P36961) and imaged on Zeiss LSM 800 with Airyscan Microscope (RRID:SCR_015963).

### Single-Cell RNA-seq Sample and Library Preparation

Dissociated single cells were fixed using the Chromium Next GEM Single Cell Fixed RNA Sample Preparation Kit (10x Genomics Cat# PN-1000414) to preserve RNA integrity. Fixed cells were then processed using the Chromium Next GEM Single Cell 3’ Reagent Kits v3.1 (10x Genomics) according to the manufacturer’s protocol. Following cDNA amplification and library construction, all libraries were pooled and sequenced on a NextSeq 2000 platform from Illumina (RRID:SCR_010233).

### Single-Cell RNA-seq Data Processing and Quality Control

Raw sequencing data were demultiplexed into FASTQ files using Cell Ranger (v7.1.0, 10x Genomics, RRID:SCR_017344) mkfastq. Sequence reads were aligned to the mm10-2020-A reference genome using Cell Ranger count, generating gene-barcode matrices. Downstream analysis was performed using the Seurat^46^ R package (v5.1.0, RRID:SCR_016341).

Gene expression matrices were loaded with Read10X_h5 funciton, and Seurat objects were created using CreateSeuratObject. For CD45+ samples, cells with >200 UMIs, >200 detected features, <20% mitochondrial gene expression, and log_10_(nFeature_RNA)/log_10_(nCount_RNA) > 0.8 were retained. For original total mixtures, the same criteria were applied, with a stricter UMI threshold (>500).

### Single-Cell RNA-seq Data Integration and Cell Type Annotation

Following quality control, datasets were normalized using NormalizeData, highly variable genes were identified using FindVariableFeatures, and data were scaled using ScaleData. Dimensionality reduction was performed using PCA, and batch correction was applied using Harmony. Clustering was conducted using FindNeighbors and FindClusters across multiple resolutions, with the optimal resolution selected using the Clustree visualization tool. UMAP plots were generated using Harmony-corrected principal components. Low-quality clusters enriched in mitochondrial genes were removed, and the remaining cells were then re-clustered. Cluster-specific markers were identified using FindAllMarkers, and the top markers were visualized to guide manual annotation. Cell type labels were assigned based on canonical marker genes and validated using SingleR^11^(RRID:SCR_023120) with the ImmGen^12^ reference dataset. To support tumor cell identification, inferCNV^13,14^ (RRID:SCR_021140) was used to infer copy number variations.

### EMT Score Calculation

To evaluate EMT states in both mouse single-cell RNA-seq and TCGA bulk RNA-seq datasets, three scoring methods were applied: the 76-gene signature (76GS) method^15,16^, the Kolmogorov–Smirnov (KS) statistic–based approach^17^, and the multiple linear regression (MLR)–based method^18^. For the mouse tumor data, EMT scores were computed at both the single-cell and pseudobulk levels using log-normalized expression values from the Seurat object’s data slot. To enable direct comparison with the TISMO^19^ bulk RNA-seq reference, pseudobulk expression values were log2-transformed and gene-wise z-score scaled. The same gene-wise scaling was applied to the TISMO bulk RNA-seq dataset. The directionality of the KS and MLR scores was inverted to align with the 76GS scoring scale for consistent interpretation. EMT scores were calculated on the combined dataset, which included 81 untreated parental murine mammary tumor samples from TISMO and the scRNA-seq–derived pseudobulk data.

### Correlation Analysis of *TACSTD2* Expression and EMT Scores

Transcriptomic and clinical data for breast cancer (TCGA-BRCA) were downloaded via the TCGAbiolinks (RRID:SCR_017683) R package. Clinical annotations were extracted from BCR XML files and processed using GDCprepare_clinic to identify samples of triple-negative breast cancer. Pearson correlation analysis was conducted to evaluate the relationship between TACSTD2 expression and EMT scores.

## Data Availability

The single-cell RNA sequencing data generated in this study will be publicly available in the Gene Expression Omnibus (GEO) under accession number GSE303505.

## Code Availability

The analysis code supporting this work is not publicly available but can be obtained from the lead author upon reasonable request.

## Author Contributions

H.L. designed and performed the experiments, conducted data analysis, interpreted the results, and wrote the manuscript. M.B. assisted with experiments and provided critical suggestions throughout the study. F.W.K. supervised single-cell RNA sequencing library preparation, sequencing, and primary data processing. L.A.S. and X.W. served as advisory committee members and provided valuable feedback on the study design, data interpretation, and manuscript preparation. B.C.C, D.R.P, and T.W.M as corresponding authors, supervised the overall project, provided guidance throughout the study, and take responsibility for the integrity of the work as a whole. All authors reviewed and approved the final manuscript.

## Acknowledgments

Preprocessing of sequencing data was carried out in the Genomics and Molecular Biology Shared Resource (GMBSR) (RRID:SCR_021293) at Dartmouth, which is supported by the National Cancer Institute (NCI) Cancer Center Support Grant 5P30CA023108 and NIH S10 award 1S10OD030242. Single-cell studies were conducted through the Dartmouth Center for Quantitative Biology in collaboration with the GMBSR, with support from the NIGMS (P20GM130454) and NIH S10 award S10OD025235.

