## Supplementary Figure S1 for "Epithelial-Mesenchymal Transition is Associated with Altered Immune Composition and Cytotoxic Function in Triple-Negative Breast Cancer"

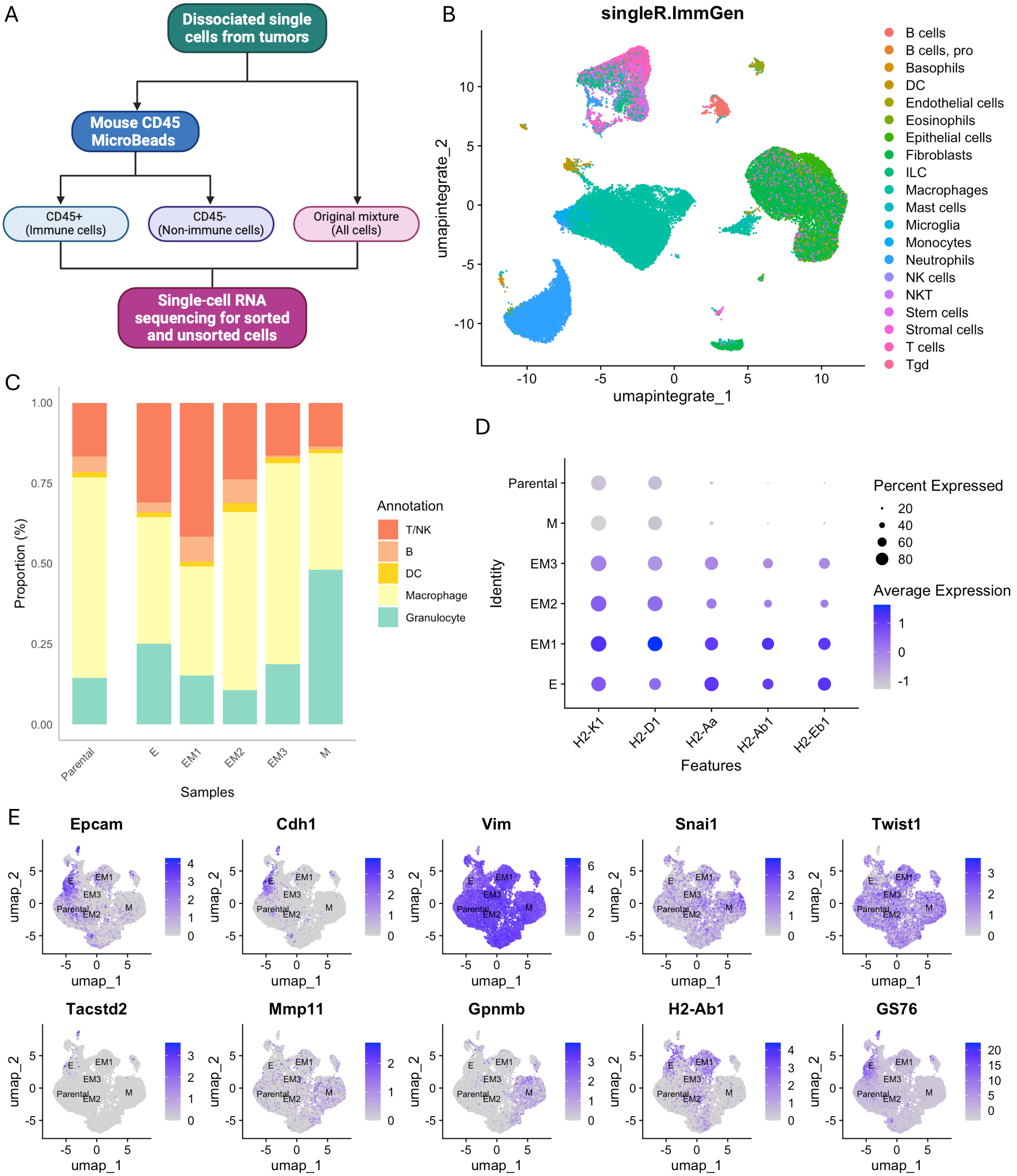
A
Dissociated single cells from tumors
Mouse CD45 MicroBeads
CD45+ (Immune cells)
CD45- (Non-immune cells)
Original mixture (All cells)
Single-cell RNA sequencing for sorted and unsorted cells
B
singleR.ImmGen
umapintegrate_2
umapintegrate_1
B cells
B cells, pro
Basophils
DC
Endothelial cells
Eosinophils
Epithelial cells
Fibroblasts
ILC
Macrophages
Mast cells
Microglia
Monocytes
Neutrophils
NK cells
NKT
Stem cells
Stromal cells
T cells
Tgd
C
Proportion (%)
Samples
Parental
E
EM1
EM2
EM3
M
Annotation
T/NK
B
DC
Macrophage
Granulocyte
D
Identity
Features
H2-K1
H2-D1
H2-Aa
H2-Ab1
H2-Eb1
Percent Expressed
Average Expression
E
Epcam
Cdh1
Vim
Snai1
Twist1
Tacstd2
Mmp11
Gpnmb
H2-Ab1
GS76
umap_1
umap_2

**Figure S1: Experimental workflow and single-cell transcriptomic analysis of cells in the tumor microenvironment.**

**A** Schematic of the experimental workflow: tumor tissues were dissociated into single cells and subjected to magnetic separation using Mouse CD45 MicroBeads to isolate CD45+ (immune) and CD45- (non-immune) cell populations. Unsorted cells were also retained to represent the original tumor cell mixture. Immune cells and the original mixture were subjected to single-cell RNA sequencing. **B** UMAP plot showing cell type annotations of scRNA-seq data from the original cell mixture, using the SingleR tool with the ImmGen reference dataset. **C** Stacked bar plot displaying the proportional distribution of major immune cell populations in original mixtures across different samples. **D** Dot plot showing the expression of MHC class I and II genes in tumor cells subsetted from the original mixture. Dot size indicates the percentage of cells expressing each gene, and color intensity reflects average expression levels. **E** UMAP feature plots showing 76GS EMT scores and expression of EMT-associated genes in tumor cells subsetted from the original mixture.
