## Supplementary Figure S2 for "Epithelial-Mesenchymal Transition is Associated with Altered Immune Composition and Cytotoxic Function in Triple-Negative Breast Cancer"

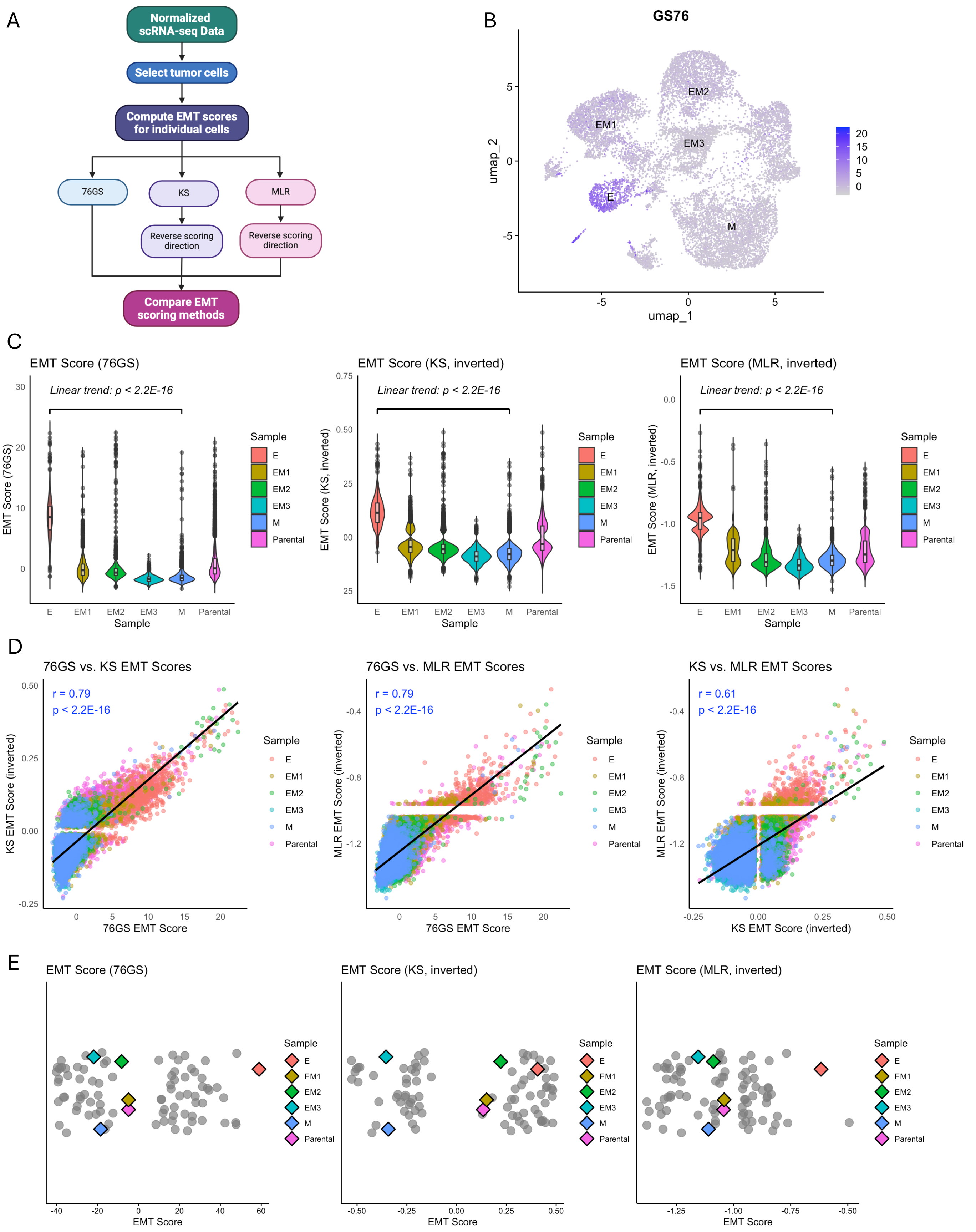

A
Normalized scRNA-seq Data
Select tumor cells
Compute EMT scores for individual cells
76GS
KS
MLR
Reverse scoring direction
Reverse scoring direction
Compare EMT scoring methods
B
GS76
EM2
EM1
EM3
E
M
umap_2
umap_1
C
EMT Score (76GS)
EMT Score (KS, inverted)
EMT Score (MLR, inverted)
Linear trend: p < 2.2E-16
Sample
E
EM1
EM2
EM3
M
Parental
D
76GS vs. KS EMT Scores
76GS vs. MLR EMT Scores
KS vs. MLR EMT Scores
r = 0.79
p < 2.2E-16
r = 0.61
76GS EMT Score
KS EMT Score (inverted)
MLR EMT Score (inverted)
E
EMT Score

**Figure S2: Comparative analysis of EMT scoring methods applied to tumor cell populations from scRNA-seq data.**

**A** Schematic workflow for calculating EMT scores at the single-cell level. Tumor cells were first identified from normalized scRNA-seq datasets. EMT scores were computed using three distinct methods: the 76-gene signature (76GS) based method, the Kolmogorov–Smirnov (KS) statistic-based method, and the multiple linear regression (MLR)-based method. For consistency across methods, the KS and MLR scores were reversed in direction to align with the 76GS scale. **B** Feature plot showing 76GS EMT scores projected onto UMAP of tumor cells, parental 4T1 cells were excluded. **C** Violin plots comparing EMT scores across samples derived from different EMT clones using the 76GS, KS (inverted), and MLR (inverted) scoring methods. Linear regression analysis was performed to assess the trend in EMT scores from E to M. **D** Pairwise scatter plots showing correlations between EMT scoring methods: 76GS vs. KS (inverted), 76GS vs. MLR (inverted), and KS vs. MLR (both inverted). **E** Comparative distribution of EMT scores between EMT clone–derived tumor cells and TISMO mammary cancer cell lines using each EMT scoring method. The same gene-wise z-score scaling was applied to both pseudobulk and TISMO datasets. Tumor cells (colored diamonds) were mapped alongside 81 untreated, parental wild-type mammary cancer cell lines (gray dots) from the TISMO database.
