## Supplementary Figure S3 for "Epithelial-Mesenchymal Transition is Associated with Altered Immune Composition and Cytotoxic Function in Triple-Negative Breast Cancer"

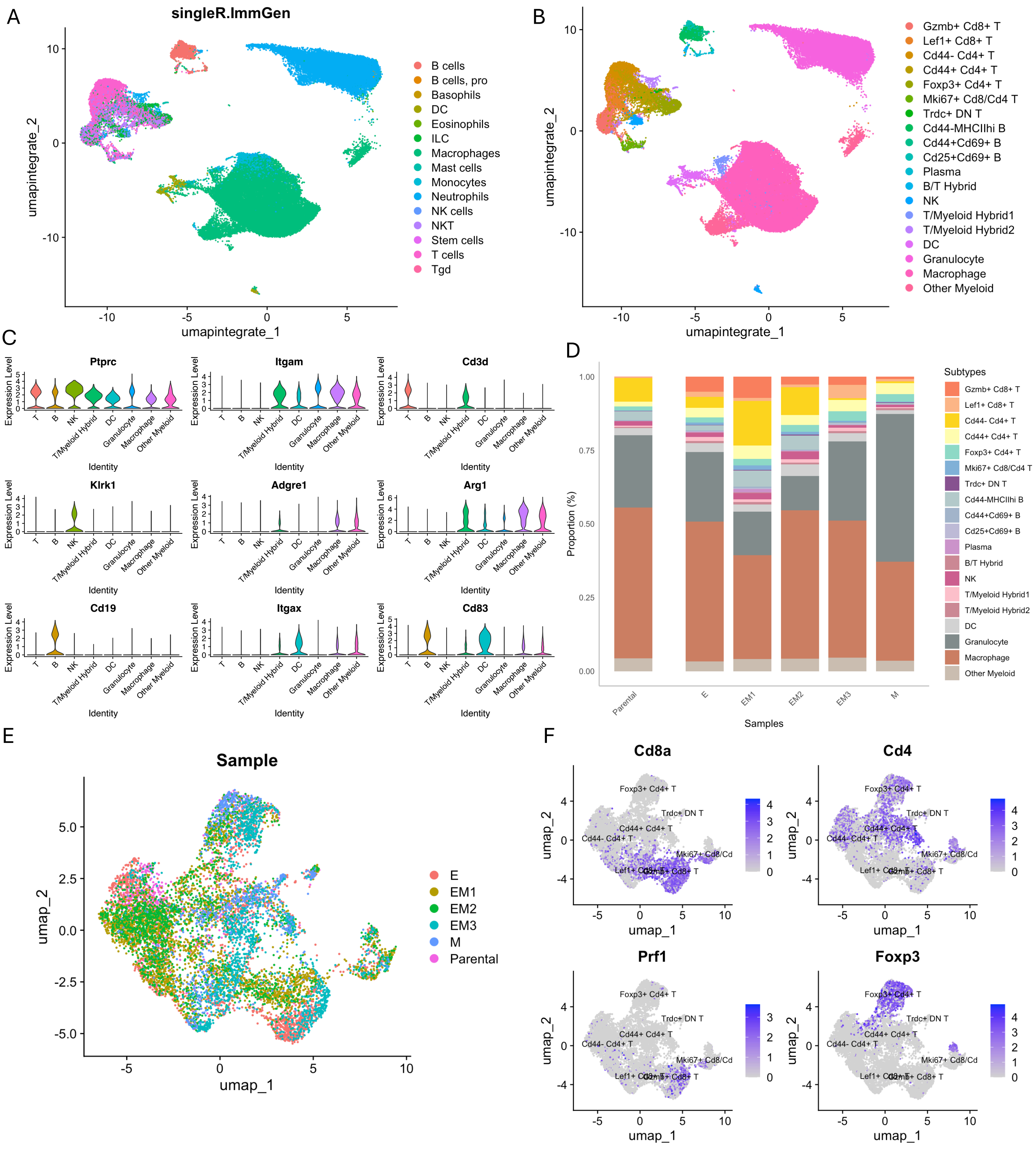
A
singleR.ImmGen
umapintegrate_2
umapintegrate_1
B cells
B cells, pro
Basophils
DC
Eosinophils
ILC
Macrophages
Mast cells
Monocytes
Neutrophils
NK cells
NKT
Stem cells
T cells
Tgd
B
Gzmb+ Cd8+ T
Lef1+ Cd8+ T
Cd44- Cd4+ T
Cd44+ Cd4+ T
Foxp3+ Cd4+ T
Mki67+ Cd8/Cd4 T
Trdc+ DN T
Cd44-MHCIIhi B
Cd44+Cd69+ B
Cd25+Cd69+ B
Plasma
B/T Hybrid
NK
T/Myeloid Hybrid1
T/Myeloid Hybrid2
DC
Granulocyte
Macrophage
Other Myeloid
C
Ptprc
Itgam
Cd3d
Klrk1
Adgre1
Arg1
Cd19
Itgax
Cd83
Expression Level
Identity
T
B
NK
T/Myeloid Hybrid
DC
Granulocyte
Macrophage
Other Myeloid
D
Proportion (%)
Samples
Parental
E
EM1
EM2
EM3
M
Subtypes
E
Sample
umap_2
umap_1
E
EM1
EM2
EM3
M
Parental
F
Cd8a
Cd4
Prf1
Foxp3

**Figure S3: Immune cell subtype characterization from CD45+ tumor-infiltrating cells.**

**A** UMAP plot showing cell type annotations of scRNA-seq data from CD45+ samples using the SingleR tool with the ImmGen reference dataset. **B** UMAP plot displaying immune cell subtypes annotations. **C** Violin plots showing the expression of canonical marker genes to identify immune cell subtypes. **D** Stacked bar plot illustrating the proportional distribution of immune cell subtypes across different CD45+ immune samples. **E** UMAP plot showing T cell distribution, colored by sample. **F** Feature plots displaying the expression of key markers genes (Cd8a, Cd4, Prf1, Foxp3) used to identify T cell subtypes.
