## Supplementary Table S1 for "Epithelial-Mesenchymal Transition is Associated with Altered Immune Composition and Cytotoxic Function in Triple-Negative Breast Cancer"

**Supplementary Table 1.** EMT-related genes

| | | | | | |
|---|---|---|---|---|---|
| ZEB1 | TNFRSF21 | EVPL | KRT19 | KDF1 | TMEM30B |
| LIX1L | TMEM45B | FXYD3 | GRHL1 | CDS1 | CLDN7 |
| VIM | MPP7 | CLDN4 | BSPRY | CHEK2 | EPCAM |
| AXL | CHAF1B | CRB3 | C1orf116 | MPZL2 | SCNN1A |
| MMP2 | SSH3 | TSKU | S100A14 | PATJ | CDH1 |
| ANTXR2 | TC2N | MAPK13 | S100A11P1 | ESRP1 | FZR1 |
| XXYLT1 | MUC1 | GALNT3 | SPINT2 | TMC4 | |
| FN1 | EPPK1 | STAP2 | ANKRD22 | ITGB6 | |
| NRP1 | SHROOM3 | DSP | ST14 | TMEM125 | |
| TGFBI | EPN3 | ELMO3 | GRHL2 | EPHA1 | |
| GALNT5 | PRSS22 | KRTCAP3 | PRR5 | ENPP5 | |
| PPARG | AP1M2 | MAL2 | TJP3 | EPB41L5 | |
| HNMT | SH3YL1 | F11R | TACSTD2 | ERBB3 | |
| CARD6 | KLC3 | ADGRF1 | CDH3 | RAB25 | |
| RBPMS | SERINC2 | ADGRG1 | CDH15 | PRSS8 | |
